## Supplemental file for "Atypical cognitive training-induced learning and brain plasticity and their relation to insistence on sameness in children with autism"

### Supplementary Materials

**I. Supplementary Methods** …. Pages 2-6.

**II. Supplementary Results** …. Pages 6-7.

**III. Supplementary Tables** …. Pages 8-15.

**IV. Supplementary Figures** …. Pages 16-21.

#### I. Supplementary Methods

##### Participants

Our pilot studies suggest a group size of 25 is sufficient for identifying greater retrieval use in ASD with power >80% and effect size Cohen's $\delta = 0.9$. To meet the goal, a total of 116 children were recruited from the San Francisco Bay Area via flyer or poster advertisements at schools, libraries, and community centers. Thirty-six children met ASD criteria based on DMS-IV and the Autism Diagnostic Interview-Revised (ADI-R) *(53)* and/or the Autism Diagnostic Observation Schedule (ADOS) *(54)*. Twenty-eight TD children included in the study had no history of genetic, neurological, psychiatric, or learning disorders, no personal and family history (first degree) of developmental disorders, and no significant difficulty during pregnancy, labor, delivery, or immediate neonatal period, or abnormal developmental milestones. All eligible participants completed the training protocol in this study. One child with ASD was excluded due to the lack of valid data in the fMRI task, with an accuracy below chance level (< 50%) and no responses to over 30% of trials in more than half of runs in the pre-training fMRI task.

##### Study design and procedure

***Overall study protocol.*** The overall study protocol is summarized in **Figure 1a**. This study consisted of the following sessions: *i*) pre-training neuropsychological (NP) assessments of cognitive abilities and clinical assessments; *ii*) pre-training task fMRI scan session (math verification task) and outside-of-scanner assessment session (math production task and strategy assessment); *iii*) one-on-one math training, in which five training days were spread out across a two-week period, with no more than three days between training days; *iv*) post-training task fMRI scan session and outside-of-scanner assessment sessions with the same tasks as pre-training.

***Training problem sets.*** All possible single-digit addition problems with operands from 2 to 9 were created, excluding ties (e.g., 2+2). Half of 28 problems had the smaller operand first, and the other half had the larger operand first. These problems were further divided into two-well balanced sets of 14 problems with the sum of each set equal to 154. Each set had 6 "non-carry over" problems with sums of 10 or less and 8 "carry over" problems with sums of 11 or more. Half of the problems in each set were randomly assigned to have double-digit in the first operand (or double-digit in the second operand), with decade values in the double-digit operands ranging from 20 to 80. The assignment of decade values to single-digit problems amongst each set of 14 problems had to follow several constraints: *i*) the numbers 2 to 9 appear at least once as the single-digit operand; *ii*) double-digit multiples of 11 are excluded; *iii*) at least one problem is summed to a value in each of the decades from 20-90; *iv*) sums are separated by at least 3 units. Finally, all sums were unique across the training sets.

***Training activities.*** On each training day, the tutor first introduced a "break-apart" (decomposition) strategy to children to facilitate their learning of complex arithmetic problem solving, after which children completed a worksheet to solve a set of trained problems using the break-apart strategy. Children were encouraged to use the memory-retrieval strategy whenever possible on the last day of training. In the 'Pirate game' task, participants played 3 rounds of a flashcard game in which problems are presented on the screen in animated soap bubbles at random order. Multiple attempts were allowed to enter the correct response before the time ran out when the bubble reached the top of the screen and the next problem appeared at the bottom of the screen. The time limit was set as 15 seconds for the first round of the Pirate Game in the first training session and was reduced by one second after each successful round (> 75% accuracy) across training days, reaching 9-11 seconds for the final round of the last training day.

Below is the list of activities that children were engaged in each day of training (Days 1 to 5). Participants accumulated stickers for completing each activity on a "treasure board" and were invited to select a small prize upon completion of 20 training activities.

***i) Warm-up flashcard.*** The first task each day, except for Day 1 of training, was a simple warm-up flashcard task, in which participants were shown each of 14 problems on a flashcard and were asked to verbally produce the solution to the problem. The tutor proceeded to the next problem once the child produced a correct answer.

***ii) Lesson and worksheet.*** After warm-up, participants were given an introductory lesson on "break-apart" (decomposition; rule-based) strategy, which involved separating the double-digit operand into a decade (multiple of 10) and a single-digit number and then adding the sum of the two single-digit numbers to the decade number (e.g. 65 + 7 => 60 + 5 + 7 = 60 + 12 = 72). On a 14-problem worksheet, the tutor demonstrated this method on the first problem, then asked the child to solve the next one with guided support. Once participants understood the strategy, they were prompted to complete the worksheet using the break-apart strategy. While participants were asked to use the break-apart strategy for the first four days of training, on Day 5, participants were encouraged to solve the problems by retrieving the answers from memory (memory-retrieval strategy).

***iii) Treasure hunt/Bingo.*** Participants played a multiple-choice game in which they were asked to match a presented solution to a problem to one of the problems on the board. There were two versions of this game, "Treasure hunt" (played on Days 1, 3, and 5) and "Bingo" (played on Days 2 and 4). In "Treasure hunt," participants were shown a treasure map containing 14 boxes with problems and a deck of 14 cards with possible solutions placed faced down. Participants drew one card at a time and placed it on one of the boxes in the treasure map to match the solution (card) to the problem (box). In "Bingo," participants received a 4 x 4 bingo card, where each square (except for two "free spots") contained a problem. Participants were given bingo chips with solutions to match to corresponding problems on the card.

***iv) Pirate game (computer).*** Participants played three rounds of a computerized flashcard game in which each of 14 problems was presented in a random order on the screen in an animated soap bubble. Participants completed each problem by typing the answer to the problem and pressing the enter key. Each bubble contained a trained problem appeared in the bottom of the screen and moved upwards until it reached the top of the screen. When the participant answered the problem correctly within the time the bubble remained on the screen, the bubble popped and the next problem appeared at the bottom of the screen. Participants were allowed as many attempts to "pop the bubble" with a correct response before time ran out and the next problem would appear on the screen. The time limit was set as 15 seconds for the first round in the first training session (Day 1) and was reduced by one second after each successful round (> 75% accuracy) across training days.

***v) Oral review.*** Participants were asked to verbally produce the answer to each of 14 problems read out loud by the tutor, without the use of worksheet or physical manipulatives. The order of problems presented varied from day to day. All participants in the same group (children assigned to training set A or B) received the same order.

***vi) Memory game.*** Participants completed a memory game using fill boards from the game "Guess Who" (Hasbro Gaming, Pawtucket, Rhode Island) boards. The 14 problems were split into two boards with seven pairs of problem and answer on each. Problems were distributed on the top two rows and solutions on the bottom two rows. All cards were initially faced down and participants were instructed to pick one problem and one solution to see if they matched. If they matched, these cards would remain face up and if they did not match, they would be turned face down and new pairs would be picked until all problems and solutions were matched. The placement of the pairs remained constant throughout the days of training and all participants completed the two memory boards in the same order. The tutor timed the duration taken to complete each board for each participant. In addition, they were encouraged to find matching problem-solution pairs in as few moves as possible to elicit memory retrieval for spatial locations.

***vii) Beat your time.*** Participants completed three rounds of timed flashcard game in which a deck of flashcards containing 14 problems was completed twice in each round. The tutor presented each card to the participant one at a time. Participants verbally produced correct answer to move onto the next problem. If an incorrect answer was produced, the tutor waited until the participant provided the correct answer before continuing. The tutor timed the duration taken to complete each round and participants were encouraged to beat the time taken to complete previous round.

***viii) Review worksheet.*** Participants completed a 14-problem worksheet at the end of every training session for review. Each day, participants were prompted to solve the problems however easiest for them. By Day 5, children were encouraged to use the memory-retrieval strategy.

**Behavioral tasks**

***Math verification task in the fMRI scanner.*** As shown in **Figure 1a**, each trial began with a fixation cross with 500ms, followed by a double-digit plus single-digit problem presented for 6 seconds. During this problem presentation phase, participants were instructed to solve the problem. Next, a possible solution to the problem (probe) was presented for up to 3 seconds. During this response phase, participants indicated whether the possible solution was same or different from the answer to the problem they were thinking of by pressing the left button with their index finger or the right button with their middle finger, after which a blank screen filled a 10-second trial length followed by a jitter period ranging from 8 to 12 seconds.

In each run, half of the probes presented were correct answers and the other half were incorrect. Within each run, correct and incorrect problems were presented in a pseudo-random order where no more than three correct or incorrect problems appeared in a row. Incorrect answers differed from the correct answer by plus or minus 1, 2, or 10. All possible differences from the incorrect sum were used once in each run, as well as one trial presented an incorrect answer of either +1 or -1 for a total of 7 incorrect answers per run. If the first run had an extra +1 trial, then the second run had the -1 trial and vice versa. Notably, answers differing by plus or minus 10 ensured that children considered both operands to solve the problems. Run were not repeated even in cases of excessive head movement, in order to ensure that all participants have the same amount of exposure to the problems. As part of quality control of data acquired, runs with an accuracy below chance level (< 50%) and no responses to 30% or more trials were considered invalid and were not included in the analysis.

***Math production task and strategy assessments.*** The problems were presented in the same order in each of pre- and post-training sessions across all children, with the order of the problems reversed between pre- and post-training sessions.

**fMRI data acquisition**

fMRI data were acquired on a 3T GE Signa scanner (General Electric, Milwaukee, WI) using a custom-built 8-channel head coil at the Richard M. Lucas Center for Imaging at Stanford University. Head movement was minimized during the scan by placing cushions around the participant’s head. During the fMRI task scanning, children held a custom-made MR-compatible computer mouse in their right hand. A total of 31 axial slices (4.0 mm thickness, 0.5 mm skip) parallel to the anterior commissure (AC)-posterior commissure (PC) line and covering the whole brain were imaged using a T2*-weighted gradient-echo spiral in-out pulse sequence *(57)* with the following parameters: repetition time (TR) = 2 seconds, echo time (TE) = 30ms, flip angle = 80°, one interleave. The field of view was 22 cm, and the matrix size was 64 × 64, providing an in-plane spatial resolution of 3.4375mm. Reduction of blurring and signal loss arising from field inhomogeneity was accomplished by the use of an automated high-order shimming method based on spiral acquisitions before fMRI acquisition *(58)*. Each task fMRI run lasted 4 minutes and 50 seconds (i.e., 145 volumes/time points) including 10 seconds at the beginning of each run for allowing scanner equilibration.

**fMRI data preprocessing**

fMRI data were analyzed with the following preprocessing procedures using SPM12 (http://www.fil.ion. ucl.ac.uk/spm/). The first 5 volumes (10 seconds) were not analyzed to allow for signal equilibration. A linear shim correction was applied separately for each slice during reconstruction based on a magnetic field map acquired automatically by the pulse sequence at the beginning of scan *(57)*. The subsequent processing included the motion correction with realigning to the first scan and the slice-timing correction. Then, images were spatially normalized to standard Montreal Neurological Institute (MNI) space using the echo-planar imaging template, resampled to 2mm isotropic voxels using trilinear sinc interpolation, and smoothed with a 6mm full-width at half-maximum Gaussian kernel. Data from 13 children with ASD and 4 TD children were excluded in the following analysis due to excessive head motion (translation or rotation above 10mm or 10° in any direction or mean frame-wise head motion above 0.5mm) in more than half of runs. Data from runs with excessive head motion were also excluded for the following analysis. Additionally, as part of quality control of acquired data, the image intensity at the pre-training session from one child with ASD was identified as an outlier

(> 3 standard deviations away from the mean) and excluded from data analysis. A gray matter mask with 172,470 voxels was generated by overlapping image mask across children and SPM gray matter template (grey matter probability > 0.2) to limit our analysis to the gray matter. Given that some participants did not have full coverage of the cerebellum and brainstem, our analysis focused on cerebral regions.

**Multivariate neural representational pattern analysis**

***Group differences.*** To examine whether children with ASD and TD children show similar or different patterns of functional brain plasticity in response to training, individual whole-brain NRP maps were submitted to two-sample *t*-test. Significant clusters were determined using a voxel-wise height threshold of $p < 0.005$ and an extent threshold of $p < 0.05$ using family-wise error correction for multiple comparisons based on Gaussian random field (GRF). Anatomical brain locations of significant clusters were identified using Harvard-Oxford atlas (65) and Juelich Histological atlas (64). Similarly, for *a priori* ROI analysis, two-sample *t*-tests were used to examine group differences in NRP. Significant results were determined by false discovery rate (FDR)-corrected $p < 0.05$. Cohen's *f* was calculated to provide estimates of effect sizes.

***Relation between NRP and learning gains.*** Next, to examine whether the relationship between functional brain plasticity and learning are similar or different between ASD and TD groups, a general linear model was used with group (ASD, TD), learning gains, and their interaction as independent variables, and NRP as the dependent variable. Here, learning gains were measured by percent changes in accuracy for trained problems in the math verification task, considering that this task administered during fMRI scanning and thus provides performance/learning related measure directly associated with brain activation/plasticity. Significant clusters were identified using a voxel-wise height threshold of $p < 0.005$ and an extent threshold of $p < 0.05$ using family-wise error correction for multiple comparisons based on GRF. For *a priori* ROI analysis, a general linear model was used to test group by learning gain interaction on NRP for each predefined region. Significant regions were determined by FDR-corrected $p < 0.05$. Cohen's *f* was calculated to provide estimates of effect sizes.

#### II. Supplementary Results

##### Learning profiles of children with ASD, compared to TD children, during training: accuracy and reaction time

To further examine whether children with ASD demonstrate comparable changes in performance during math training relative to TD children, follow-up analyses were performed for accuracy and reaction time separately. For accuracy, a 5x2 (Session x Group) repeated measures ANOVA showed a significant main effect of session ($F(4, 212) = 8.21$, $p < 0.001$, $\eta^2_p = 0.13$), but no main effect of group or session by group interaction ($F$s $\leq 1.01$, $p$s $\geq 0.405$). For reaction time, a 5x2 (Session x Group) repeated measures ANOVA showed a significant main effect of session ($F(4, 212) = 127.81$, $p < 0.001$, $\eta^2_p = 0.71$), but no main effect of group or session by group interaction ($F$s $\leq 1.42$, $p$s $\geq 0.230$). These findings indicate that children with ASD improve as well as their TD children during training on both accuracy and reaction time.

**Changes in problem-solving strategy use in response to training in children with ASD, compared to TD children: strategy consistency**

To estimate how consistent the dominant strategy is in each participant, we calculated the dominant strategy rate by dividing the number of problems using the dominant strategy by all possible problems (i.e., 14 trained problems). No significant group differences on dominant strategy rate were found on either before (two-sample *t*-test, $t(59) = 0.41$, $p = 0.683$; ASD: M = 0.67, SD = 0.21; TD: M = 0.65, SD = 0.18) or after training (two-sample *t*-test, $t(59) = -0.48$, $p = 0.630$; ASD: M = 0.76, SD = 0.21; TD: M = 0.78, SD = 0.20).

**Changes in problem-solving strategy use in response to training in children with ASD, compared to TD children: untrained problems**

To further address the specificity of differential changes in dominant strategy with training between groups for trained problems, we examined children's changes in dominant strategy for untrained problems. Children with ASD and TD children showed a comparable distribution of dominant strategy use for untrained problems before and after training (all $\chi^2 \leqslant 0.58$, $p \geqslant 0.444$) (Supplementary Figure 2b and Supplementary Table 5), confirming the specificity of group differences in problem-solving strategy use for trained problems in response to training.

**Training-related neural representational plasticity in children with ASD, compared to TD children: joint contribution of MTL and IPS**

To further address explore the degree of joint contribution of MTL and IPS regions, key brain regions typically associated with math learning, we performed a multiple regression model with NRP of bilateral MTL and right IPS identified from the whole brain analysis as predictors. Here, MTL and IPS jointly contributed to learning gain in each group, ASD: adjusted $R^2 = 0.37$; $F(3, 17) = 4.96$; $p = 0.012$; TD: adjusted $R^2 = 0.31$; $F(3, 20) = 4.48$; $p = 0.015$, which indicates the importance of both brain regions.

**Training-related neural representational plasticity in children with ASD, compared to TD children: untrained problems**

To investigate the specificity of brain-behavior relationship in ASD and TD groups observed for trained problems, we examined interaction between group and performance changes for untrained problems in the brain regions identified from the whole brain analysis (group by learning gain interaction) for trained problems and a priori defined regions related to math learning. No significant effect was observed for untrained problems in these brain regions (**Supplementary Figures 5-6**).

**Influence of insistence on sameness on the relationship between training-related brain plasticity and learning in children with ASD: control regions**

To address whether the mediation findings are specific MTL and IPS, we examined V1 and the whole brain. Specifically, we defined V1 as the 17 area from Brodmann and used the grey matter template from the main analysis. No significant effect was observed for V1 ($b = -0.22$, $se = 0.25$, *t* = -0.86, *p* = 0.406) or the whole brain (*b* = -0.29, *se* = 0.32, *t* = -0.93, *p* = 0.370) in children with ASD.

#### III. Supplementary Tables

##### Supplementary Table 1. Demographic and clinical measures

| Measure | ASD (n = 35) | TD (n = 28) | $t/\chi^2$ | *p* |
|---|---|---|---|---|
| **Gender (M/F)** | 29/6 | 22/6 | 0.19[a] | 0.667[a] |
| **Age (years)** | 9.98 ± 0.92 | 10.00 ± 1.09 | -0.06 | 0.953 |
| **WASI scale** | | | | |
| Verbal IQ | 112.54 ± 14.33 | 119.14 ± 13.01 | -1.89 | 0.063 |
| Performance IQ | 119.40 ± 19.44 | 114.07 ± 11.23 | 1.29 | 0.203 |
| Full IQ | 117.71 ± 15.72 | 118.64 ± 9.41 | -0.28 | 0.784 |
| **ADI-R** | | | | |
| Social | 20.09 ± 6.14 | | | |
| Verbal | 16.17 ± 4.46 | | | |
| Repetitive behavior | 5.43 ± 2.83 | | | |
| Development | 3.17 ± 1.07 | | | |
| **ADOS** | | | | |
| Social/Affect | 9.26 ± 3.00[†] | | | |
| Restricted and repetitive behavior | 2.91 ± 1.48[†] | | | |
| Severity scores | 7.18 ± 1.78[†] | | | |
| Total | 12.18 ± 3.62[†] | | | |
| **RRIB sub-scores (based on ADI)** | | | | |
| Insistence on sameness | 1.10 ± 1.30[††] | | | |
| Circumscribed interests | 2.62 ± 1.52[††] | | | |
| Repetitive motor behavior | 2.75 ± 1.72[††] | | | |

The mean, standard deviation of measure, *t* or $\chi^2$, and *P* value of two sample t-test/chi-square test are shown here. Abbreviations: ASD, children with autism spectrum disorder; TD, typically developing children; M, male; F, female; IQ, intelligence quotient; WASI, Wechsler Abbreviated Scale of Intelligence; ADI-R, Autism Diagnostic Interview-Revised (diagnostic scores); ADOS, Autism Diagnostic Observation Schedule-new algorithm; RRIB, restricted and repetitive interests and behaviors.

[a] The statistic value were obtained using a chi-square test.

[†] Data from one participant are missing.

[††] Data from two participants are missing.

**Supplementary Table 2. Number of participants included in each analysis**

| | ASD | TD |
|---|---|---|
| **Behavioral analysis** | | |
| Training task | 29 | 26 |
| Math verification task | 35 | 28 |
| Math production task and strategy assessment | 33 | 28 |
| **Brain measures and brain-behavior analysis** | 21 | 24 |
| **RRIB-brain-behavior moderation analysis** | 19 | n.a. |

Abbreviations: ASD, children with autism spectrum disorder; TD, typically developing children; n.a., not available.

**Supplementary Table 3. Results of repeated measures ANOVA for behavioral performance**

| | Measure | Effect | *F* | *df* | $\eta^2_p$ | BF | *p* |
|---|---|---|---|---|---|---|---|
| **Training task** | **ES** | Time Effect | 113.15 | 4,212 | **0.68** | **>100** | **<0.001** |
| | | Group Effect | 0.04 | 1,212 | <0.01 | 0.41 | 0.849 |
| | | Interaction | 1.13 | 4,212 | 0.02 | 0.14 | 0.344 |
| | | **ASD** | | | | | |
| | | Time Effect | 70.46 | 4,112 | **0.72** | **1.56** | **<0.001** |
| | | **TD** | | | | | |
| | | Time Effect | 45.02 | 4,100 | **0.64** | **1.90** | **<0.001** |
| **Math verification task** | **ACC** | **Trained** | | | | | |
| | | Time Effect | 9.09 | 1,61 | **0.13** | **8.11** | **0.004** |
| | | Group Effect | 1.60 | 1,61 | 0.03 | 0.59 | 0.210 |
| | | Interaction | 0.48 | 1,61 | 0.01 | 0.31 | 0.492 |
| | | **Untrained** | | | | | |
| | | Time Effect | 1.31 | 1,61 | 0.02 | 0.29 | 0.257 |
| | | Group Effect | 0.57 | 1,61 | <0.01 | 0.40 | 0.453 |
| | | Interaction | 1.90 | 1,61 | 0.03 | 0.56 | 0.173 |
| **Math production task** | **RT** | **Trained** | | | | | |
| | | Time Effect | 148.13 | 1,59 | **0.72** | **>100** | **< 0.001** |
| | | Group Effect | 2.87 | 1,59 | 0.05 | 0.87 | 0.096 |
| | | Interaction | 1.65 | 1,59 | 0.03 | 0.65 | 0.204 |
| | | **Untrained** | | | | | |
| | | Time Effect | 13.72 | 1,59 | **0.19** | **61.54** | **<0.001** |
| | | Group Effect | 2.20 | 1,59 | 0.04 | 0.85 | 0.144 |
| | | Interaction | 6.06 | 1,59 | **0.09** | **3.59** | **0.017** |

#### Supplementary Table 4. Results of *t*-tests for behavioral performance

| Task | Measure | Contrast | Mean ± SD | | *t* | df | Cohen's *d* | BF | *p* |
|---|---|---|---|---|---|---|---|---|---|
| **Training task** | **ES** | **ASD vs. TD** | ASD | TD | | | | | |
| | | Session 1 | 0.17±0.07 | 0.18±0.08 | -0.53 | 53 | -0.14 | 0.46 | 0.60 |
| | | Session 2 | 0.22±0.1 | 0.21±0.09 | 0.29 | 53 | 0.08 | 0.29 | 0.78 |
| | | Session 3 | 0.25±0.11 | 0.24±0.10 | 0.21 | 53 | 0.06 | 0.30 | 0.84 |
| | | Session 4 | 0.27±0.11 | 0.26±0.11 | 0.30 | 53 | 0.08 | 0.29 | 0.77 |
| | | Session 5 | 0.30±0.11 | 0.28±0.12 | 0.44 | 53 | 0.12 | 0.28 | 0.66 |
| | **Learning rate** | **ASD vs. TD** | ASD | TD | | | | | |
| | | | 0.03±0.01 | 0.03±0.02 | 1.28 | 53 | 0.35 | 0.27 | 0.206 |
| **Math verification task** | **ACC -trained** | **Pre vs. Post** | Pre | Post | | | | | |
| | | ASD | 0.86±0.11 | 0.88±0.10 | -1.72 | 34 | -0.29 | 0.68 | 0.095 |
| | | TD | 0.88±0.11 | 0.92±0.06 | -2.54 | 27 | **-0.48** | **2.91** | **0.017** |
| | | **ASD vs. TD** | ASD | TD | | | | | |
| | | Pre | 0.86±0.11 | 0.88±0.11 | -0.73 | 61 | -0.19 | 0.32 | 0.467 |
| | | Post | 0.88±0.10 | 0.92±0.06 | -1.67 | 61 | -0.42 | 0.83 | 0.101 |
| | **ACC gain -trained** | **ASD vs. TD** | ASD | TD | | | | | |
| | | | 0.04±0.11 | 0.06±0.12 | -0.73 | 61 | -0.18 | 0.32 | 0.470 |
| | **ACC -untrained** | **Pre vs. Post** | Pre | Post | | | | | |
| | | ASD | 0.84±0.12 | 0.84±0.13 | -0.17 | 34 | -0.03 | 0.74 | 0.864 |
| | | TD | 0.88±0.11 | 0.84±0.13 | 1.73 | 27 | 0.33 | 0.18 | 0.095 |
| | | **ASD vs. TD** | ASD | TD | | | | | |
| | | Pre | 0.84±0.12 | 0.88±0.11 | -1.40 | 61 | -0.35 | 0.59 | 0.168 |
| | | Post | 0.84±0.13 | 0.84±0.13 | -0.02 | 61 | 0.00 | 0.26 | 0.988 |
| | **ACC gain -untrained** | **ASD vs. TD** | ASD | TD | | | | | |
| | | | 0.02±0.16 | -0.04±0.14 | 1.35 | 61 | 0.34 | 0.55 | 0.183 |
| **Math production task** | **RT (s) -trained** | **Pre vs. Post** | Pre | Post | | | | | |
| | | ASD | 6.53±2.89 | 3.85±2.21 | 8.68 | 32 | **1.51** | **0.17** | **<0.001** |
| | | TD | 5.31±2.28 | 3.15±1.69 | 9.31 | 27 | **1.76** | **0.17** | **<0.001** |
| | | **ASD vs. TD** | ASD | TD | | | | | |
| | | Pre | 6.53±2.89 | 5.31±2.28 | 1.80 | 59 | 0.46 | 1.00 | 0.078 |
| | | Post | 3.85±2.21 | 3.15±1.69 | 1.37 | 59 | 0.35 | 0.57 | 0.178 |
| | **RT gain -trained** | **ASD vs. TD** | ASD | TD | | | | | |
| | | | -0.41±0.19 | -0.41±0.16 | -0.03 | 59 | -0.01 | 0.28 | 0.979 |
| | **RT (s) -untrained** | **Pre vs. Post** | Pre | Post | | | | | |
| | | ASD | 6.77±2.91 | 5.47±2.91 | 3.70 | 32 | **0.64** | **38.68** | **<0.001** |
| | | TD | 5.26±2.40 | 5.00±2.59 | 1.35 | 27 | 0.26 | 0.46 | 0.187 |
| | | **ASD vs. TD** | ASD | TD | | | | | |
| | | Pre | 6.77±2.91 | 5.26±2.40 | 2.19 | 59 | **0.56** | **1.88** | **0.033** |
| | | Post | 5.47±2.91 | 5.00±2.59 | 0.66 | 59 | 0.17 | 0.31 | 0.512 |
| | **RT gain -untrained** | **ASD vs. TD** | ASD | TD | | | | | |
| | | | -0.19±0.24 | -0.06±0.19 | -2.23 | 59 | **-0.57** | **1.57** | **0.029** |

#### Supplementary Table 5. Results of chi-squared tests for dominant strategy use

| | | | | Memory-based | Rule-based | $\chi^2$ | *df* | $\phi$ | BF | *p* |
|---|---|---|---|---|---|---|---|---|---|---|
| **Dominant strategy use** | **Group vs. Strategy (trained)** | **Pre** | **ASD** | 4 | 29 | 0.40 | 1 | 0.08 | 0.50 | 0.312 |
| | | | **TD** | 5 | 23 | | | | | |
| | | **Post** | **ASD** | 17 | 16 | 4.81 | 1 | **0.28** | **3.39** | **0.028** |
| | | | **TD** | 22 | 6 | | | | | |
| | **Group vs. Strategy (untrained)** | **Pre** | **ASD** | 4 | 29 | 0.03 | 1 | 0.02 | 0.46 | 0.864 |
| | | | **TD** | 3 | 25 | | | | | |
| | | **Post** | **ASD** | 11 | 22 | 0.58 | 1 | 0.10 | 0.42 | 0.444 |
| | | | **TD** | 12 | 16 | | | | | |
| **Strategy differentiation between trained and untrained problems** | **Problem vs. Strategy (ASD)** | **Pre** | **Trained** | 4 | 29 | 0 | 1 | 0.00 | 0.43 | 1.000 |
| | | | **Untrained** | 4 | 29 | | | | | |
| | | **Post** | **Trained** | 17 | 16 | 2.23 | 1 | 0.18 | 0.90 | 0.135 |
| | | | **Untrained** | 11 | 22 | | | | | |
| | **Problem vs. Strategy (TD)** | **Pre** | **Trained** | 5 | 23 | 0.58 | 1 | 0.10 | 0.57 | 0.445 |
| | | | **Untrained** | 3 | 25 | | | | | |
| | | **Post** | **Trained** | 22 | 6 | 7.49 | 1 | **0.37** | **13.24** | **0.006** |
| | | | **Untrained** | 12 | 16 | | | | | |

**Supplementary Table 6. Brain regions showing significant group by behavior interaction on neural representational plasticity between pre- and post-training for trained problems**

| Region | MNI coordinates | | | Peak ($F$) | Cluster size (voxel) | Cohen's $f^2$ |
|---|---|---|---|---|---|---|
| | x | y | z | | | |
| **R MTL/PHG** | 30 | -48 | 2 | 28.43 | 211 | 0.69 |
| **L MTL/PHG** | -16 | -36 | -14 | 23.97 | 108 | 0.58 |
| **R FEF/preCG** | 28 | -20 | 60 | 21.27 | 209 | 0.52 |
| **R IPS/LOC** | 30 | -64 | 50 | 20.54 | 116 | 0.50 |
| **R LOC** | 30 | -78 | 16 | 19.59 | 76 | 0.48 |
| **R MFG/FP** | 28 | 38 | 28 | 18.55 | 95 | 0.45 |

These regions showed significant group by behavior interaction on NRP for trained problems at the threshold of $p < 0.005$ height and $p < 0.05$ cluster extent, GRF corrected. The learning gains, computed as changes in accuracy for trained problems in the verification task during the fMRI scan, was used as behavioral measure in this analysis. L, Left; R, Right; MTL, medial temporal lope; PHG, Parahippocampal gyrus; FEF, Frontal eye fields; preCG, Precentral gyrus; IPS, Intraparietal sulcus; LOC, Lateral occipital cortex; MFG, Middle frontal gyrus; FP, Frontal pole; NRP, neural representational plasticity.

**Supplementary Table 7. Results of brain-behavior association between region of interest (ROI)-based neural representational plasticity and learning gains.**

| Region | $F$ | df | Cohen's $f^2$ | $p$ |
|---|---|---|---|---|
| **L MTL** | 8.15 | 1,41 | **0.20** | **0.007** |
| **R MTL** | 8.87 | 1,41 | **0.22** | **0.005** |
| **L IPS** | 8.08 | 1,41 | **0.20** | **0.007** |
| **R IPS** | 0.06 | 1,41 | <0.01 | 0.805 |

The group by behavior interaction effects on NRP for trained problems at the threshold of $p < 0.05$ FDR corrected. The learning gains was computed as changes in accuracy for trained problems in the verification task during the fMRI scan. L, Left; R, Right; MTL, Medial temporal lope; IPS, Intraparietal sulcus; NRP, Neural representational plasticity.

**Supplementary Table 8**. **Moderation results for RRIB sub-scores on the association between brain and behavioral measures.**

| | RRIB sub-scores | Interaction | | | | Model | | |
|---|---|---|---|---|---|---|---|---|
| | | ***b*** | ***se*** | ***t*** | ***p*** | $R^2$ | ***F*** | ***p*** |
| **NRP (R MTL) - learning gains** | Insistence on sameness | -0.09 | 0.29 | -0.32 | 0.755 | 0.33 | 2.50 | 0.099 |
| | Circumscribed interests | 0.36 | 0.32 | 1.12 | 0.281 | 0.28 | 1.97 | 0.162 |
| | Repetitive motor behavior | -0.48 | 0.38 | -1.28 | 0.221 | 0.34 | 2.60 | 0.091 |
| **NRP (L MTL) - learning gains** | Insistence on sameness | **-0.85** | **0.31** | **-2.76** | **0.015** | **0.53** | **5.64** | **0.009** |
| | Circumscribed interests | -0.21 | 0.26 | -0.80 | 0.437 | 0.33 | 2.51 | 0.098 |
| | Repetitive motor behavior | 0.04 | 0.40 | 0.11 | 0.916 | 0.31 | 2.25 | 0.125 |
| **NRP (R IPS) - learning gains** | Insistence on sameness | **-0.44** | **0.19** | **-2.28** | **0.038** | **0.34** | **4.09** | **0.026** |
| | Circumscribed interests | 0.01 | 0.32 | 0.03 | 0.975 | 0.07 | 1.46 | 0.264 |
| | Repetitive motor behavior | -0.09 | 0.28 | -0.34 | 0.738 | 0.08 | 1.52 | 0.250 |

RRIB, Repetitive and restricted interests and behaviors; NRP, Neural representational plasticity; MTL, Medial temporal lope; IPS, Intraparietal sulcus.

#### IV. Supplementary Figures

| Set A | Set B |
|---|---|
| 2 + 84 | 2 + 27 |
| 8 + 36 | 28 + 4 |
| 38 + 3 | 3 + 62 |
| 87 + 4 | 34 + 6 |
| 75 +8 | 46 +5 |
| 9 + 47 | 5 + 53 |
| 3 + 56 | 52 + 9 |
| 25 + 2 | 6 + 37 |
| 6 + 29 | 69 + 3 |
| 6 + 72 | 7 + 48 |
| 4 + 65 | 73 + 4 |
| 57 + 5 | 8 + 82 |
| 64 + 9 | 85 + 9 |
| 43 + 7 | 9 + 78 |

**Supplementary Figure 1.** Problem sets used in Set A and Set B. Problem sets were balanced in structure and task difficulty (see **Methods** for details).

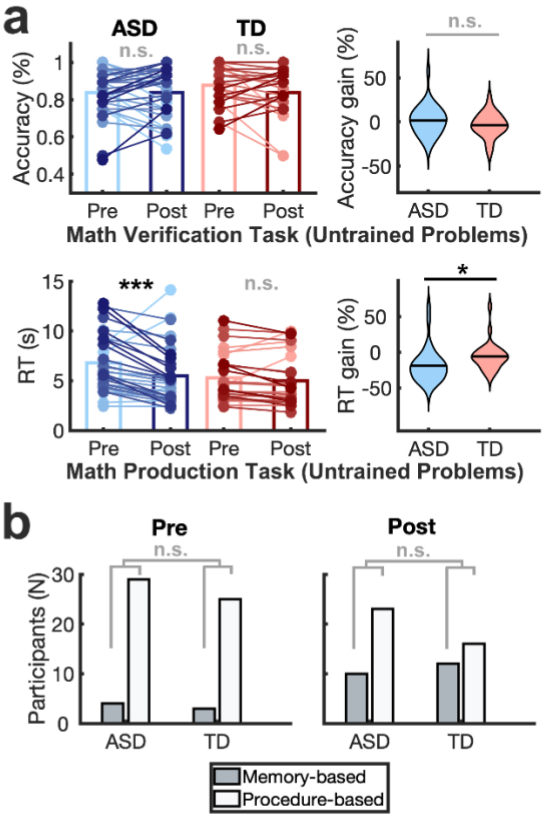


**Supplementary Figure 2. Changes in performance and strategy use for untrained problems in children with autism spectrum disorder (ASD) and typically developing (TD) children in response to training. a. Changes in performance.** Accuracy for untrained problems of math verification task in the fMRI scanner at pre- and post-training is shown for each group (*top*). Accuracy gain, measured as percentage of change from pre- to post-training ($Accuracy_{gain}$ = ($Accuracy_{post}$ - $Accuracy_{pre}$)/$Accuracy_{pre}$), was not significantly different between groups. Reaction time (RT) for untrained problems of math production task outside the scanner at pre- and post-training is shown for each group (*Bottom*). RT gain, measured as percentage of change from pre- to post-training ($RT_{gain}$ = ($RT_{post}$ - $RT_{pre}$)/$RT_{pre}$), was significantly different between groups, with children with ASD becoming faster to solve the problems correctly with training while TD children did not significantly change for untrained problems. The pair of dots connected by a line represents the performance of the same child at pre- and post-training, with darker blue/red dots/lines indicating greater gains in performance. The bars indicate the mean value in each group at pre- and post-training (lighter and darker blue/pink). **b. Changes in dominant strategy use for untrained problems.** The distribution of participants using memory-based and rule-based strategies as their dominant strategy for untrained problems before and after training is shown for each group. Children with ASD showed similar distribution of dominant strategy use as TD children for untrained problems at pre- and post-training. *** $p < 0.001$; ** $p < 0.01$; * $p < 0.05$; *n.s.*, not significant.

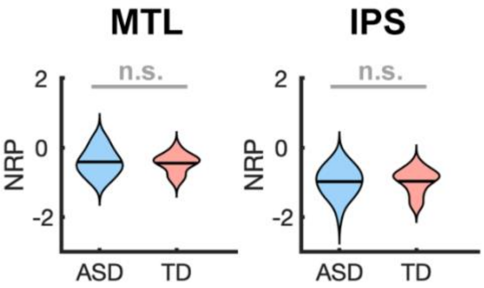


**Supplementary Figure 3. Group differences in neural representation plasticity (NRP) for right hemisphere regions of interest (ROIs).** Mean NRP across individuals in the right medial temporal lobe (MTL) and right intraparietal sulcus (IPS) was comparable between children with ASD and TD children. n.s., not significant.

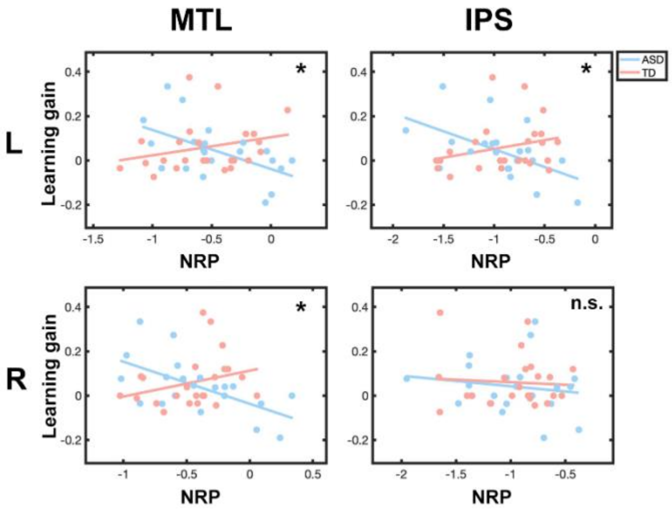


**Supplementary Figure 4. Results of brain-behavior association between region of interest (ROI)-based neural representational plasticity (NRP) and learning gains for trained problems in children with autism spectrum disorder (ASD), compared to typically developing (TD) children.** ROI-based analysis showed significant interaction between group and learning gain on NRP for trained problems in bilateral medial temporal lobe (MTL) and left intraparietal sulcus (IPS). L, Left; R, Right; *, $p < 0.05$, FDR-corrected; n.s., not significant.

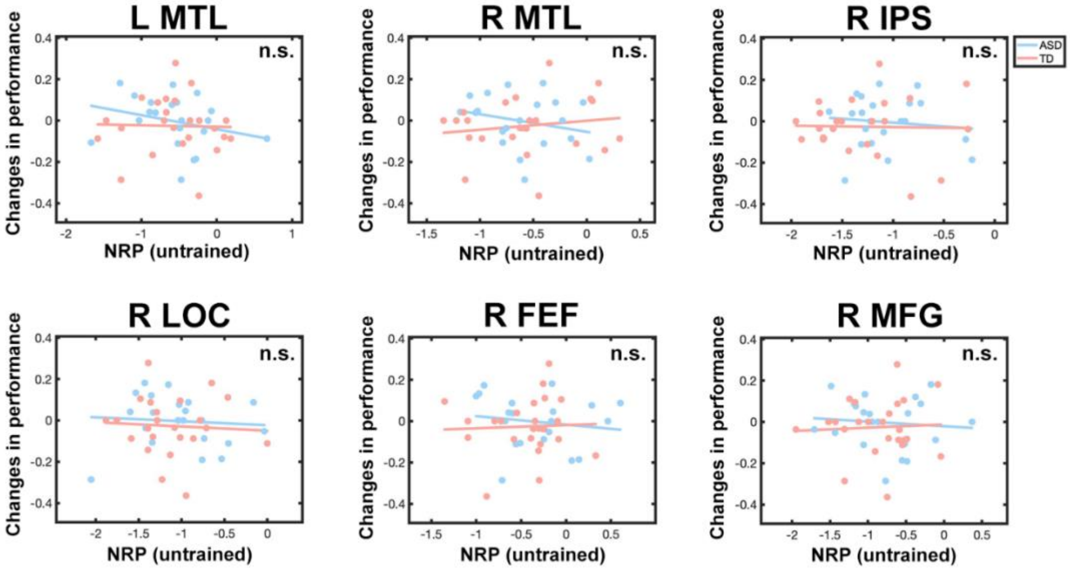


**Supplementary Figure 5. Brain-behavior association between neural representational plasticity (NRP) and changes in performance for untrained problems in children with autism spectrum disorder (ASD) relative to typically developing (TD) children.** Brain regions identified from whole brain analysis for trained problems did not show significant interaction between group and change in performance for untrained problems. L, Left; R, Right; MTL, medial temporal lobe; IPS, intraparietal sulcus; LOC, lateral occipital cortex; FEF, frontal eye field; MFG, middle frontal gyrus; NRP, neural representational plasticity; n.s., not significant.

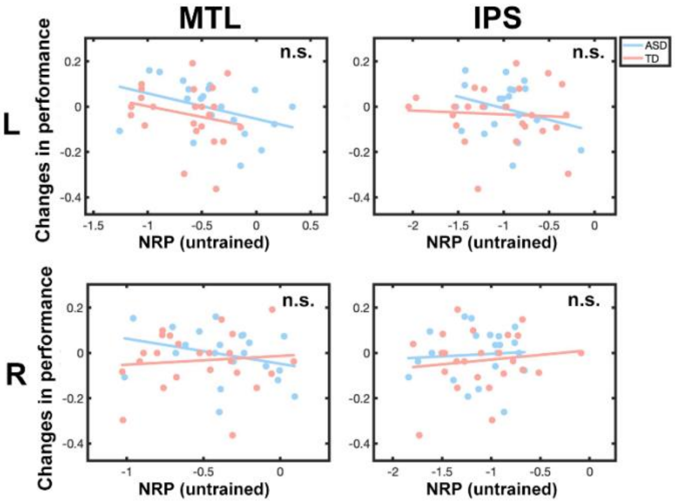


**Supplementary Figure 6. Brain-behavior association between region of interest (ROI)-based neural representational plasticity (NRP) and changes in performance for untrained problems in children with autism spectrum disorder (ASD) relative to typically developing (TD) children.** ROI-based analysis did not show significant interaction between group and changes in performance for untrained problems in regions related to math learning, including medial temporal lobe (MTL) and intraparietal sulcus (IPS). L, Left; R, Right; n.s., not significant.
